# Development of high throughput method for the analysis of anthelmintic resistance allele frequencies in field populations of gastrointestinal nematodes

**DOI:** 10.1101/569863

**Authors:** Neil D. Sargison, Madison MacLeay, Alison A. Morrison, David J. Bartley, Mike Evans, Umer Chaudhry

**Affiliations:** University of Edinburgh Roslin Institute and Royal (Dick) School of Veterinary Studies, Easter Bush Veterinary Centre, Roslin, Midlothian, EH25 9RG, United Kingdom; Moredun Research Institute, Pentlands Science Park, Bush Loan, Penicuik EH26 0PZ, United Kingdom

**Keywords:** Gastrointestinal nematode, *Teladorsagia circumcincta*, anthelmintic drug, benzimidazole, isotype 1 β-tubulin locus, deep amplicon sequencing

## Abstract

Drug resistant helminths have become a major cause of poor health and production in sheep and goats, and there is a need for diagnostic markers and tools to determine the frequency of resistance alleles in field parasite populations. Gastrointestinal nematode resistance to benzimidazole drugs is caused by a mutation in one of three positions on the isotype 1 β-tubulin locus, and in the absence of markers for resistance to other broad spectrum anthelmintic classes, these provide a relevant study example. Determination of the prevalence of these single nucleotide polymorphisms in field gastrointestinal nematode populations can be impractical using conventional molecular methods, which may be error prone or lack sensitivity at low levels of resistance. Here, we report the development of a novel method based on an Illumina Mi-seq deep amplicon sequencing platform; to sequence the isotype 1 β-tubulin locus of the small ruminant gastrointestinal nematode, *Teladorsagia circumcincta*, and determine the frequency of the benzimidazole resistance mutations. We validated the method by assessing sequence representation bias in the isotype 1 β-tubulin locus, comparing the results of Illumina Mi-seq and pyrosequencing, and applying the method to populations containing known proportions of resistant and susceptible L_3_. Finally, we applied the method to field samples collected from ewes and lambs on over a period of one year on three farms, each highlighting different aspects of sheep management and approaches to parasite control. The results show opportunities to build hypotheses with reference to selection pressures leading to differences in resistance allele frequencies between sampling dates, farms and ewes or lambs, and to consider the impact of their genetic fixation or otherwise. This study provides proof of concept of a practical, accurate, sensitive and scalable method to determine frequency of anthelmintic drug resistance mutations in gastrointestinal nematodes in field studies and as a management tool for livestock farmers.

## 1. Introduction

Resistance to broad spectrum anthelmintic drugs is a global threat to efficient livestock production (Kaplan, 2004; Kaplan and Vidyashankar, 2012). Anthelmintic drugs belonging to the tubulin binding benzimidazole class (Borgers et al., 1975) are the mainstay of gastrointestinal nematode control in lower and middle income countries throughout Africa and Asia. The benzimidazole drugs are still routinely used in intensive small ruminant production throughout Europe, despite the widespread emergence and gene flow of resistance to the drug group in most small ruminant production limiting gastrointestinal nematode species (Borgsteede et al., 1991; Fleming et al., 2006). A large survey of Scottish sheep farms showed the benzimidazole resistance phenotype in 64% of farms, almost exclusively in the predominant gastrointestinal nematode species, *Teldorsagia circumcincta* (Bartley et al., 2003). Benzimidazole resistance is an emerging problem around the world in regions where efficient small ruminant production is essential in poverty alleviation (Lalljee et al., 2018) and where alternative anthelmintic drug groups are hitherto unavailable or prohibitively expensive (Sargison et al., 2017; Van Wyk et al., 1989; Wong and Sargison, 2018).

Benzimidazole resistance is associated with mutations in the isotype 1 β-tubulin gene that prevent drug binding (Geary et al., 1992). A single nucleotide polymorphism (SNP) mutation was first identified in benzimidazole resistant *Haemonchus contortus* and *Trichostrongylus colubriformis* resulting in an amino acid substitution from tyrosine to phenylalanine at position 200 (F200Y) of the polypeptide encoded by the isotype 1 β-tubulin gene (Kwa et al., 1994; Kwa et al., 1995). The same point mutation was also shown in benzimidazole resistant *T. circumcincta* (Elard et al., 1996). This F200Y SNP is considered to be the most important mutation conferring benzimidazole resistance, and has been shown to be functionally significant with respect to the benzimidazole resistance phenotype by transfection of the gene into *Caenohabditis elegans* (Kwa et al., 1995). A substitution from tyrosine to phenylalanine at position 167 (F167Y) of the polypeptide encoded by the isotype 1 β-tubulin gene has been found in *T. circumcincta* and *H. contortus* (Silvestre and Cabaret, 2002). A third isotype 1 β-tubulin mutation (E198A) with substitution from glutamate to alanine at position 198 was first identified in field populations of *H. contortus* (Chaudhry et al., 2015; Ghisi et al., 2007; Rufener et al., 2009) and has subsequently been reported in *T. circumcincta* with two substitutions (E198L [<u>TT</u>A] or E198A [G<u>C</u>A]) (Redman et al., 2015).

The emergence and spread of benzimidazole resistance in parasite populations is aided by high effective population sizes and high mutation rates (Chaudhry et al., 2015; Redman et al., 2015). Selection for anthelmintic resistance is influenced by the timing and frequency of treatments with reference to the proportion of the total gastrointestinal nematode population that is exposed to the drug, or in a refuge from exposure, referred to as *in refugia* (van Wyk, 2001). Pragmatic advice on reducing the selection pressure for anthelmintic resistance is generally based around avoidance of selection of pre-existing resistant nematodes by affording them a competitive advantage over susceptible nematodes. Another major influence is the movement of animals infected with drug resistant parasites, leading to gene flow (Kaplan and Vidyashankar, 2012; Redman et al., 2015; Skuce et al., 2010). Anthelmintic resistance selection and gene flow pressures vary within and between seasons and with changing management; hence in the absence of conclusive evidence in support of any single risk factor, the most pragmatic advice given is to instigate mitigation strategies in response to monitoring the resistance status of individual flocks or herds (Abbott et al., 2012).

*In vivo* methods for the diagnosis of benzimidazole resistance, such as the faecal egg count reduction test (FECRT), confirm anthelmintic efficacy by comparing the number of eggs shed in host faeces before and after treatment (Coles et al., 1992; Coles et al., 2006). However, the FECRT for benzimidazole efficacy requires a sometimes impractical period of 14 days between each measurement and has poor sensitivity, in particular when less than 25% of the parasite population is resistant (Martin et al., 1989). *In vitro* bioassays, such as the egg hatch test (EHT), expose parasites to titrated concentrations thiabendazole and measure the number of first stage larvae that hatch. However, the EHT depends on sometimes impractical extraction of freshly voided eggs and also has poor sensitivity at low levels of resistance (Taylor et al., 2002).

Various molecular assays and platforms, including conventional PCR (Kwa et al., 1994), combined use of PCR with a restriction enzyme (Lehrer et al., 1995), allele specific PCR (Elard et al., 1999), cloning and Sanger sequencing analysis (Ghisi et al., 2007), real-time PCR (Álvarez-Sánchez et al., 2005), single strand conformation polymorphism (SSCP) genotyping (Skuce et al., 2010), and pyrosequence genotyping (Samson-Himmelstjerna and Blackhall, 2005), have been used to identify the isotype 1 β-tubulin SNPs in gastrointestinal nematodes. Of these, only real-time PCR and pyrosequencing can provide a practical estimate of allele frequency from pooled samples (Samson-Himmelstjerna et al., 2007), as opposed to the more laborious, hence impractical for field studies, genotyping of large numbers of individual larvae. However, while concordance between the results of these methods for pooled samples is high (Samson-Himmelstjerna et al., 2009), real-time PCR and pyrosequencing are imprecise in detecting low benzimidazole resistance allele frequencies of less than 10% (Walsh et al., 2007) and 5% (Demeler et al., 2013; Esteban-Ballesteros et al., 2017), respectively, which may be important in the study of field populations.

Pyrosequencing requires a new primer for each base pair, hence is restricted to short fragment (7 - 10 bp) sequencing. Deep amplicon sequencing using Illumina Mi-seq can accurately sequence up to about 400 bp reads with 0.1% prevalence when used to study nematode species compositions of mixed populations (Avramenko *et al*., 2015). The method might, therefore, be more sensitive in detecting low frequencies of resistance mutations than conventional methods; while allowing investigation of all three SNPs involved in benzimidazole resistance at once. Of the available next generation sequencing platforms, Illumina Mi-seq is least error prone, and best suited to high throughput. In the present study, we report the use of deep amplicon sequencing using Illumina Mi-seq for the detection of benzimidazole resistance in *T. circumcincta*. A series of experiments was performed to develop and validate the Illumina Mi-seq method for measuring resistance allele frequency. First, the number of PCR cycles to amplify DNA before sequencing was varied to identify any sequence representation bias. Second, the results from Illumina Mi-seq of *T. circumcincta* laboratory populations were compared to those from pyrosequencing assays. Third, pools were made from resistance allele genotyped individual third stage larvae (L_3_) gDNA to validate the Illumina Mi-seq assay. Finally, the assay was applied to *T. circumcincta* field populations to accurately identify the frequencies of benzimidazole resistance mutations.

## 2 Materials and methods

### 2.1. Parasite materials

Experimentally passaged and stored *T. circumcincta* laboratory populations were obtained from the Moredun Research Institute. Similar numbers of (L_3_) came from six populations: MTci2, MTci5, MTci7, MTci11, MTci12, and MTci13. The MTci2 Weybridge population was first isolated before the advent of broad spectrum anthelmintic drugs and had been phenotypically characterised as being susceptible to benzimidazole drugs (Bartley et al., 2015). The MTci5 population had originated from a sub-flock of sheep kept in fields that were formerly part of Farm 2 in the current study and had been phenotypically characterised as being resistant to benzimidazole, levamizole and ivermectin drugs (Sargison et al., 2001; Bartley et al., 2004; 2005). The MTci7 population had originated from a terminal sire sheep flock in south-east Scotland where moxidectin had been over-used in lambs following the diagnosis of resistance to other anthelmintic drug classes (Wilson and Sargison, 2007), and had been phenotypically characterised as being benzimidazole, levamizole, ivermectin and moxidectin resistant. MTci11 had been selected from MTci2 by monepantel treatment and was reported to be benzimidazole susceptible (Bartley et al., 2015). MTci12 and MTci13 had been selected from MTci7 and MTci5, respectively, by monepantel treatment and were reported to be phenotypically benzimidazole resistant (Bartley et al., 2015).

Pools of about 200 L_3_ were created, by taking 50 µl aliquots from corresponding dilutions in H_2_0 of each of the six laboratory populations. Three replicate pools of random number of larvae (∼ 200 L_3_) of each *T. circumcincta* laboratory population were used with 25, 30, 35, or 40 first round Illumina Mi-seq PCR cycles to assess accuracy and determine any sequence representation bias. Three replicate pools of random number of larvae (∼200 L_3_) of each *T. circumcincta* laboratory population were used for the comparison of the Illumina Mi-seq with the pyrosequencing assays.

To quantitatively validate the identification of benzimidazole susceptible or resistant L_3_, 48 individual MTci2 L_3_ and 48 individual MTci12 L_3_ were first picked into H_2_0. Lysates were then prepared for pyrosequence genotyping of the isotype 1 β-tubulin codon 200 locus. 1ul of lysate from known genotyped individual L_3_ was used to create three replicates each of five admixtures of homozygous susceptible (S: T<u>T</u>C) and homozygous resistant (R: T<u>A</u>C) L_3_ gDNA.

*T. circumcincta* field populations were derived from ewes and lambs on three neighbouring farms in the south-east of Scotland over multiple time points during 2016 and 2017. Farm 1 was a lowground farm with an open sheep flock of about 370 crossbred; Farm 2 was a lowground farm with an open sheep flock of about 680 crossbred ewes; Farm 3 was an extensive hill farm with a closed sheep flock of about 700 hefted Scottish Blackface ewes. There was no movement of animals, or shared grazing between Farm 1 and Farms 2 or 3, but occasional straying between Farms 2 and 3 would have been possible. The faecal samples were collected per rectum, or freshly voided onto the pasture. Ethical approval was acquired through Veterinary Ethics Review Committee (VERC) at the University of Edinburgh, reference number VERC 10 16 and consent was given by the farms’ managers. Following faecal trichostrongyle nematode egg counting using a salt flotation method with a potential sensitivity of one egg per gram (Christie and Jackson, 1982), coprocultures were set up for the recovery of third stage larvae (MAFF, 1986).

### 2.2. Genomic DNA Extraction

gDNA was prepared using 1,000 μl Direct PCR lysis reagent (Viagen), 50 μl proteinase K solution (Qiagen), and 50μl 1M dithiothreitol (DTT). To extract gDNA from individual L_3_, 25 μl of the mixture was added to each well of a 96-well plate before transferring the L_3_. The plate was placed on a thermocycler to incubate at 60°C for 2 hours to lyse cells followed by 85°C for 15 minutes to inactivate the proteinase K. For pooled L_3_, the samples were first centrifuged for 2 minutes at 7,200 xg. The supernatant was discarded, and the pellet re-suspended in 50 μl of lysis buffer, which was then placed on a thermocycler with the same conditions described for individual larvae preparation. Genomic DNA was stored at −80°C for later use.

### 2.3. Illumina Mi-seq deep amplicon sequencing of the T. circumcincta isotype 1 β-tubulin locus

Illumina Mi-seq was used to sequence a 276 bp fragment of isotype 1 β-tubulin spanning the F200Y, F167Y, and E198L or E198A SNPs. In the first round PCR, a 276 bp fragment of isotype 1 β-tubulin of *T. circumcincta* was amplified with four forward and four reverse adaptor primers (Supplementary Table S1). The PCR reaction contained 0.5 μl of KAPA HiFi polymerase (KAPA Biosystems), 0.75 μl dNTPs (10 mM), 5 μl 5X KAPA HiFi Fidelity buffer (KAPA Biosystems), 0.75 μl of each for forward and reverse primer mix (10 mM), 13.25 μl nuclease-free water, and 1 μl template DNA. The thermocycling conditions were 95°C for 2 minutes, 35 cycles of 98°C for 20 seconds, 60°C for 15 seconds, 72°C for 15 seconds, then a final extension of 72°C for 2 minutes and hold at 10°C. PCR products were purified with AMPure XP Magnetic Beads (1X) (Beckman coulter, Inc.) using a special magnetic stand (DynaMag) in accordance with the protocols described by Beckman coulter, Inc.

The second round PCR was performed by using 16 forward and 24 reverse barcoded primers. (Supplementary Table S2). Each sample was amplified using a unique combination of barcode primers. The PCR reaction contained 0.5 μl of KAPA HiFi polymerase (KAPA Biosystems), 0.75 μl dNTPs (10 mM), 5μl 5X KAPA HiFi Fidelity buffer (KAPA Biosystems), 1.25 μl of each primer (10 mM), 13.25 μl nuclease-free water and 2 μl of the first round PCR product as a template. The thermocycling conditions of the PCR were 98°C for 45 seconds, followed by 7 cycles of 98°C for 20 seconds, 63°C for 20 seconds, and 72°C for 2 minutes. PCR products were purified with AMPure XP Magnetic Beads (1X) according to the protocols described by Beckman Coulter, Inc.

The pooled library was measured with KAPA qPCR library quantification kit (KAPA Biosystems, USA). The prepared library was then run on an Illumina MiSeq sequencer using a 500-cycle pair end reagent kit (MiSeq Reagent Kits v2, MS-103-2003) at a concentration of 15 nM with addition 25% Phix Control v3 (Illumina, FC-11-2003). The MiSeq separated all sequences by sample during post-run processing by recognised indices and to generate FASTQ files.

### 2.4. Illumina Mi-seq data handling and statistical analysis

Illumina Mi-seq data analysis was performed with a bespoke pipeline using Mothur v1.39.5 software (Schloss et al., 2009) and the Illumina Mi-seq SOP (Kozich et al., 2013). In summary, raw paired-end reads were made into contigs, and too long reads (>350 bp) or ambiguous bases were removed. The reads were collapsed into a list of unique sequences. Consensus sequences of *T. circumcincta* isotype 1 β-tubulin were downloaded from the NCBI database and aligned by MUSCLE using Geneious v10.2.5 software and trimmed to the region amplified by the primers. The unique sequences were aligned with the consensus sequences and trimmed. For *T. circumcincta* laboratory populations, sequences that occurred in 1 and 2 reads were removed due to low frequency, which were most likely due to sequencing error rather than real polymorphism. The sequences derived from the field populations were more variable and sequences with less than 1000 reads were removed. The frequency of benzimidazole resistance mutations was calculated by dividing the number of sequences reads that contained the mutation by the total number of reads in each sample.

The effect of PCR cycle number on resistance frequency was analysed by running a Kruskal-Wallis rank sum test for each population. Statistical significance in this test would imply a sequence representation bias. A chi-square test was used to determine whether there was a significant difference between allele frequencies for the comparison of the samples with same proportions of benzimidazole resistant and susceptible individual *T. circumcincta* L_3_ and for the comparison of the Illumina Mi-Seq deep amplicon sequencing with the pyrosequencing assay.

### 2.5. Pyrosequence genotyping of the T. circumcincta isotype 1 β-tubulin locus

A 276 bp fragment of isotype-1 β-tubulin spanning F200Y, F167Y, and E198L or E198A was amplified with the BTUB_FOR and biotin labelled BTUB_REV primers previously described by Silvestre and Humbert (2002) and Skuce et al. (2010) (Supplementary Table S1). The PCR reaction contained 0.5 μl of 5 U/μl Taq polymerase (New England Biolabs), 0.5 μl dNTPs (10 mM), 5 μl 10X standard buffer (New England Biolabs), 0.5 μl of each primer (10 mM), 42 μl nuclease-free water and 1μl DNA template. The thermocycling parameters were 95°C for 5 minutes, 35 cycles of 95°C for 1 minute, 60°C for 1 minute, and 72°C for 1 minute, then a final extension at 72°C for 15 minutes and hold at 10°C. Following PCR amplification, the relative frequency of the F200Y, F167Y, and E198L or E198A SNPs were determined by separate pyrosequencing primers using the PyroMark ID system using the allele quantification (AQ) mode of the PSQ 96 single nucleotide position software (Biotage, Sweden). Previously, pyrosequencing primers were successfully used to genotype the *T. circumcincta* isotype 1 β-tubulin locus from individual larvae using SNP mode (Skuce et al., 2010) with some technical limitations to run the pooled samples, therefore, new pyrosequencing primers were designed to target each mutation in a pooled populations (Supplementary Table S2). PyroMark Q96 ID reagents (Qiagen) were used according to the user manual. The base dispensation was set to GTAGCTGTC for codon 200, GTAGCTCA for codon 167, and GATATCGA for codon 198. Peak heights of samples with an unknown number of larvae or individual larvae were measured either in AQ or SNP mode of the PSQ 96 single nucleotide position software previously described by Chaudhry et al. (2015).

## 3 Results

### 3.1. The effect of first round PCR cycle number on the frequency of the isotype 1 β-tubulin locus SNPs in larval pools

To assess accuracy and determine any sequence representation bias, three replicates each of mock pools were amplified from six *T. circumcincta* laboratory populations using four different numbers of first round PCR cycles. According to a Kruskal-Wallis rank sum test (performed separately for each of the mock populations), differences in the proportion of resistant or susceptible alleles at each SNP with the number of PCR cycles were not statistically significant (overall *H*_(3)_<0.2, *p*>0.9). Therefore, no sequence representation bias was observed in the benzimidazole resistance allele frequencies of the laboratory populations. The F200Y (T<u>A</u>C) SNP was found at different frequencies of between 67.2% and 94.5% in the four phenotypically benzimidazole resistant MTci5, MTci7 MTci12 and MTCi13 populations and consistently between 3% and 11.6% in the two phenotypically benzimidazole susceptible MTci2 and MTci11 populations (Fig. 1 and Supplementary Table S3). The benzimidazole resistance-associated F167Y (T<u>A</u>C) SNP was identified at a frequency of between 2% and 4.3% in the MTci13 population (Fig. 1 and Supplementary Table S3). The benzimidazole resistance associated E198L (<u>TT</u>A) and E198A (G<u>C</u>A) SNPs were not detected in any of the populations. Subsequent work used 30 cycles for the first round PCR.

**Fig. 1:**
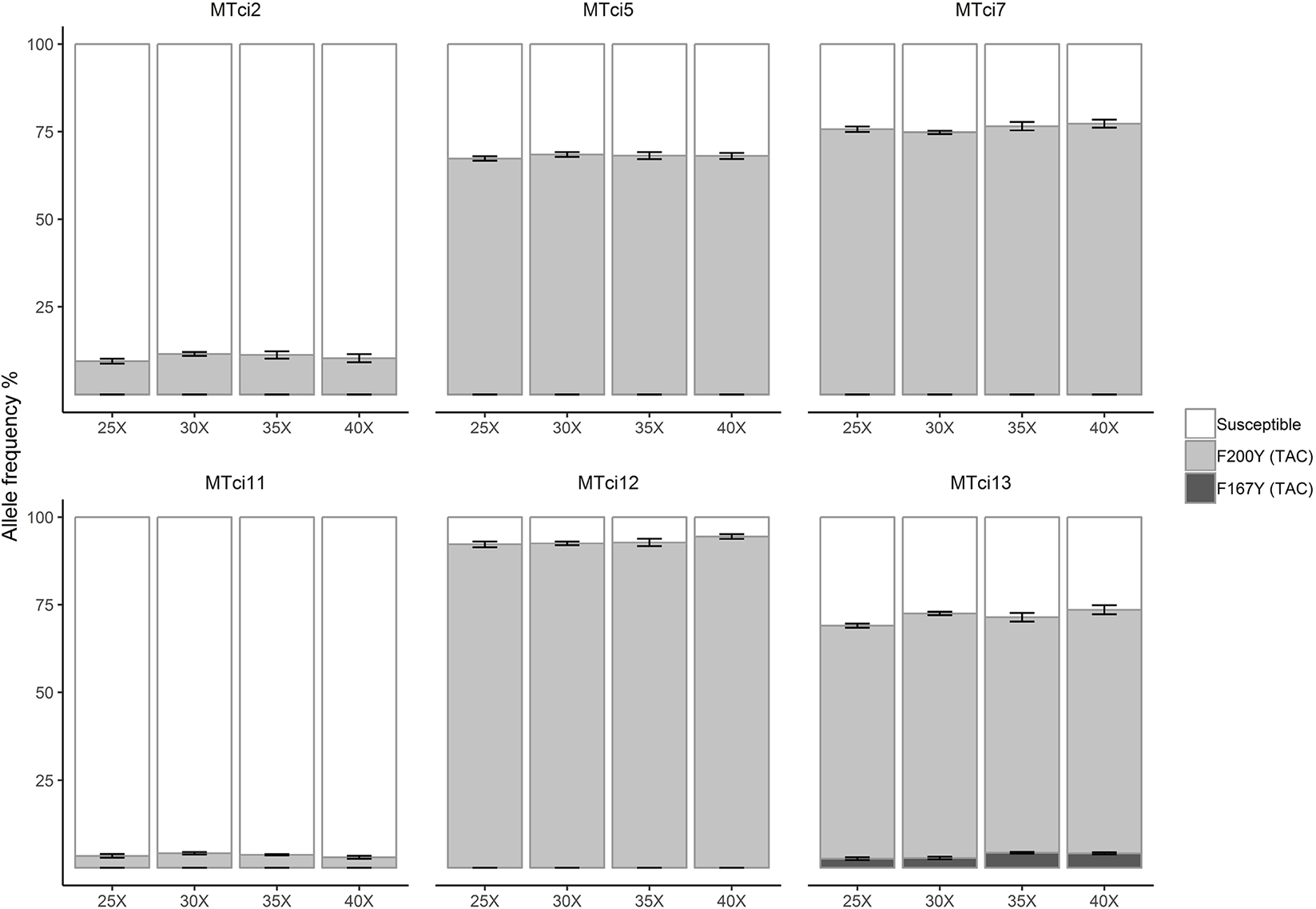
Average frequency of isotype 1 β-tubulin locus SNPs using four different PCR cycles for six *T. circumcincta* laboratory populations. The F200Y (T<u>A</u>C) resistance alleles were identified in all six populations with different frequencies. The F167Y (T<u>A</u>C) was found only in the MTci13 population (Supplementary Table S3). Dark grey shade indicates F167Y (T<u>A</u>C), medium grey indicates F200Y (T<u>A</u>C) and white indicates susceptible alleles. X-axis represents the four PCR cycles numbers (25X, 30X, 35X, 40X) and Y-axis represents the allele frequencies. Error bars represent the standard error of the mean.

### 3.2. Comparison of the Illumina Mi-Seq deep amplicon sequencing and pyrosequencing assays to determine the frequency of benzimidazole resistance SNPs

For the comparison of the Illumina Mi-seq with the pyrosequencing assay, three replicates each of mock pools were taken from six *T. circumcincta* laboratory populations. While there were variations between the results of the two methods (with the Illumina MiSeq method apparently more sensitive at low SNP frequencies), differences in the frequency of the benzimidazole resistance SNPs determined by Illumina Mi-seq and pyrosequencing were not statistically significant (Chi-square test: χ^2^ = 3.962, *p*=0.4112). The F200Y (T<u>A</u>C) SNP was found at different frequencies in the different populations of between 11.3% and 94.5% in the Illumina Mi-seq and 8.2% and 85.3% in the pyrosequencing assay (Fig. 2 and Supplementary Table S4). The F200Y (T<u>A</u>C) and F167Y (T<u>A</u>C) SNPs were only detected at low frequencies of 3.7% and 4.5% by Illumina MiSeq in the MTci11 and MTci13 populations, respectively, but were not identified by pyrosequencing assay (Fig. 2 and Supplementary Table S4). The benzimidazole resistance associated E198L (<u>TT</u>A) and E198A (G<u>C</u>A) SNPs were not detected by either method in any of the populations.

### 3.3. Validation of the proportions of benzimidazole resistant and susceptible individual T. circumcincta L_3_

To assess the accuracy of the Illumina Mi-seq platform in identifying the frequencies of phenotypically benzimidazole susceptible MTci2 and resistant MTci12 L_3_, individual L_3_ were first genotyped by pyrosequencing for the F200Y mutation. This identified homozygous susceptible (S: T<u>T</u>C) and homozygous resistant (R: T<u>A</u>C) alleles. Three replicates taken from MTci2 and MTci12 admixtures were created of individual L_3_ gDNA that had been pyrosequence genotyped, allowing gDNA pools to be produced with known allele frequencies at the isotype 1 β-tubulin codon 200 locus and used to validate the Illumina MiSeq method. There was no statistically significant difference between the expected and observed frequencies of susceptible and resistance alleles in a chi-square test, which suggests that the Illumina MiSeq method was accurate in measuring resistance allele frequencies (Chi-square test; mixS: χ^2^_(1)_<0.001, *p*=1; mixR: exact match; mixSR: χ^2^_(1)_<0.001, *p*=1; mixSRR: χ^2^_(1)_=0.057, *p*=0.845; mixSSR: χ^2^_(1)_=0.065, *p*=0.799) (Fig. 3, Supplementary Table S5). For the pools of 100% susceptible or resistance alleles (MixS(F200Y-T<u>T</u>C)-100, MixR(F200Y-T<u>A</u>C)-100], the expected and observed results were perfectly matched (Fig. 3). For pools of 33%, 50% or 67% susceptible and 33%, 50% or 67% resistance alleles [MixSR(F200Y-T<u>T</u>C/T<u>A</u>C)-50/50, MixSRR(F200Y-T<u>T</u>C/T<u>A</u>C)-25/75 MixSSR(F200Y-T<u>T</u>C/T<u>A</u>C)-75/25], the expected and observed results were nearly accurate, with not-significant variations between replicates (Fig. 3 and Supplementary Table S5).

**Fig. 2:**
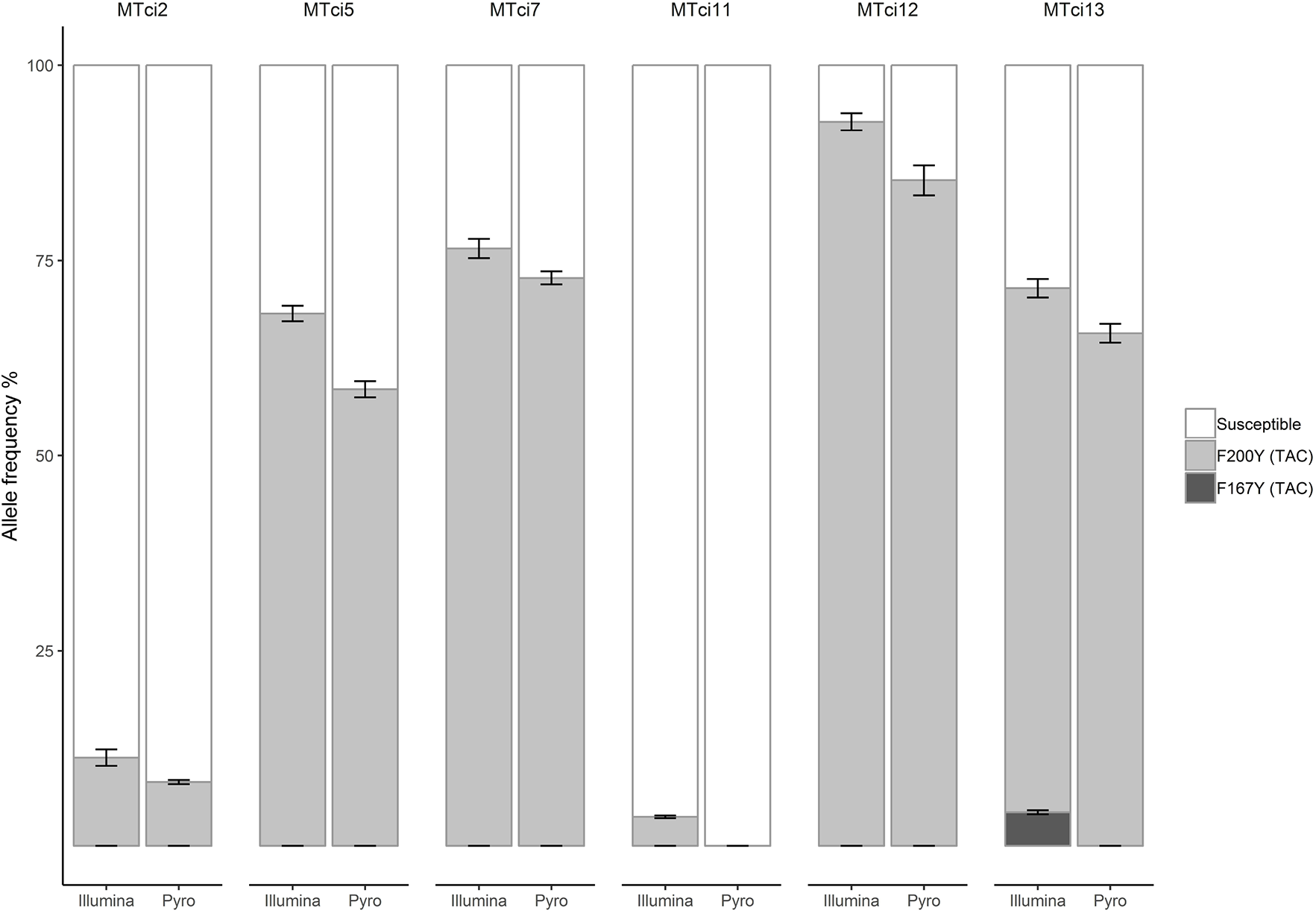
Average frequency of isotype 1 β-tubulin locus SNPs in six *T. circumcincta* laboratory populations, determined by Illumina Mi-seq and pyrosequencing. Dark grey shading indicates the F167Y (T<u>A</u>C) SNP, medium grey shading indicates the F200Y (T<u>A</u>C) SNP and white indicates susceptible alleles. Error bars represent the standard error of the mean.

**Fig. 3:**
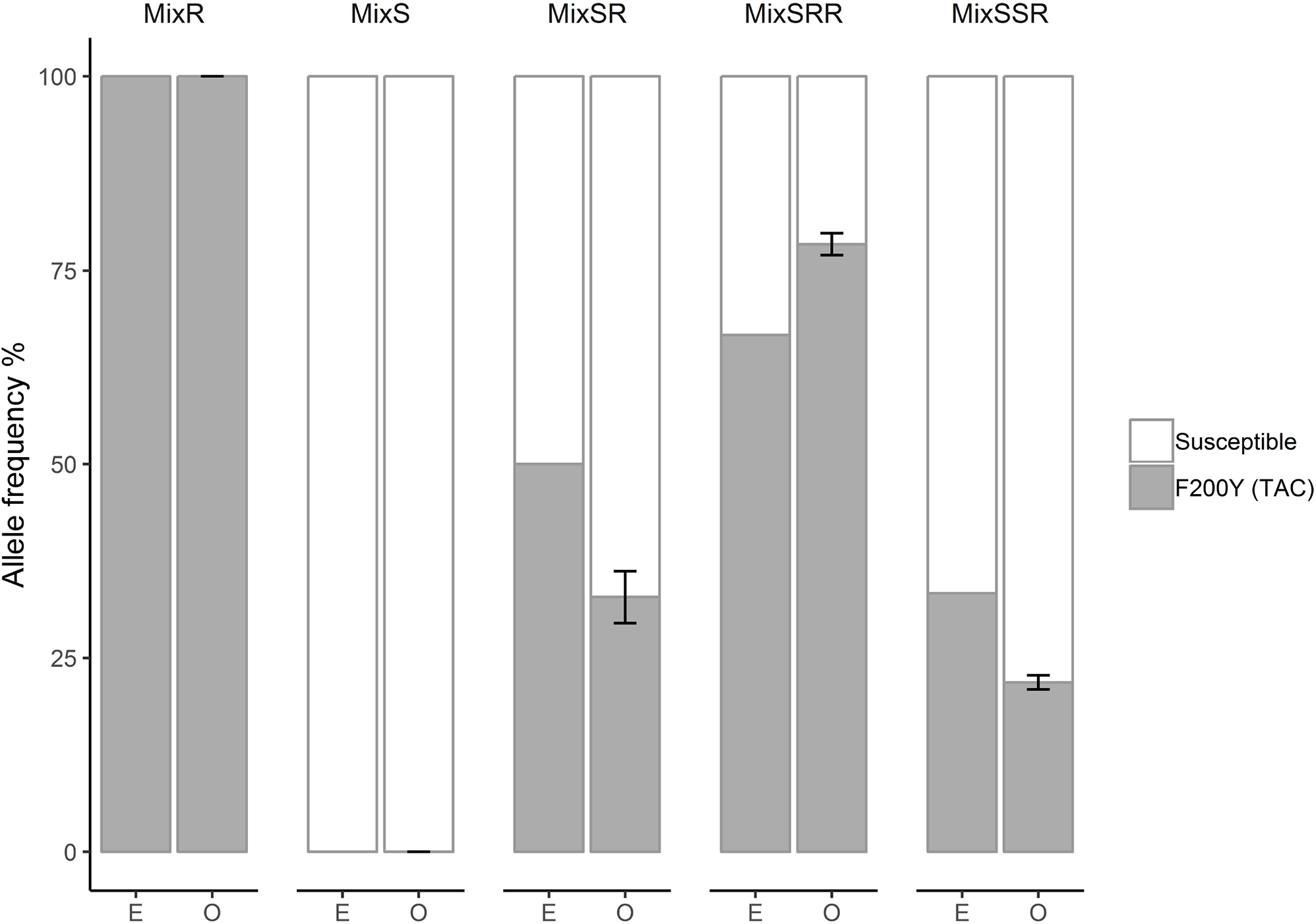
Average frequency of the isotype 1 β-tubulin locus F200Y (T<u>A</u>C) SNP in mock pools was made from different mixing of pyrosequence genotyped MTci2 individual homozygous susceptible and MTci12 homozygous resistant *T. circumcincta* L_3_. The symbol mix represents the mixing of resistant-R and susceptible-S alleles. In the X-axis, E represents the expected allele frequencies based on pyrosequence genotyping and O represents the observed allele frequencies based on Illumina MiSeq.

### 3.4. Detection of isotype 1 β-tubulin locus SNPs in T. circumcincta field populations using the Illumina Mi-Seq deep amplicon sequencing method

The Illumina Mi-Seq deep amplicon sequencing results were considered to be sufficiently consistent with isotype 1 β-tubulin pyrosequence genotyping for the platform to be evaluated on field samples. In total, field samples of 43 ewes and 31 lambs from three farms were collected at multiple time points in 2016 and 2017 (Supplementary Table S6). The allele frequencies of benzimidazole resistance mutations are shown in Fig. 4. For Farm 1, the F200Y resistance allele frequency was between 78% and 100% in both ewes and lambs. For Farm 2, the F200Y resistance allele frequency was between 61.7% and 80.2% in the lambs, but varied between 10% and 92% in the ewes. For Farm 3, the F200Y resistance allele frequency was between 64.6% and 90.2% in the lambs, but varied between 25.3% and 88.5% in the ewes. The F167Y allele frequency was between 0.5% and 6.6% on all three farms. The E198L (T<u>T</u>A) allele frequency was between 0.1% and 13.9% on all three farms.

**Fig. 4:**
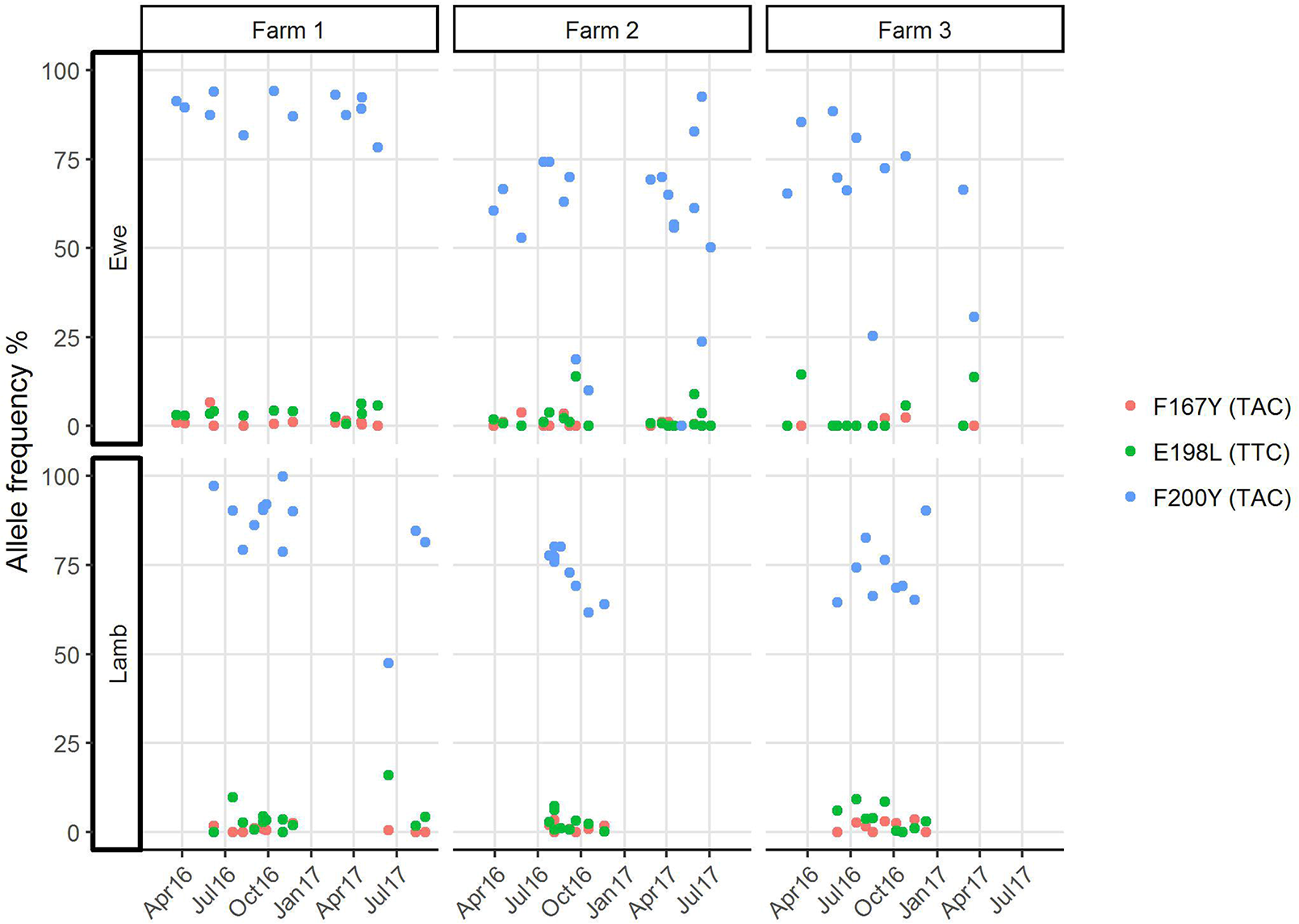
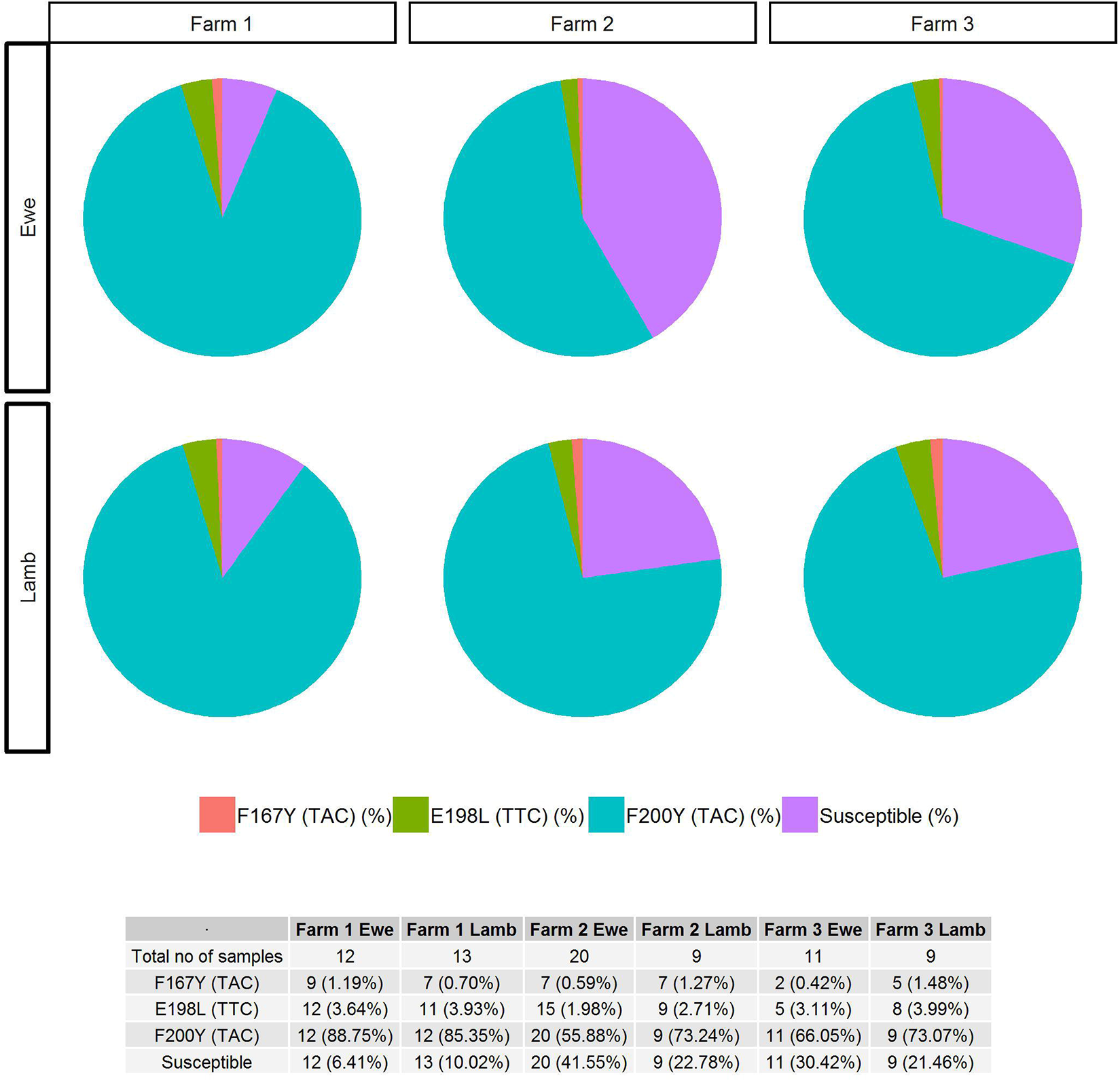
Allele frequencies of isotype 1 β-tubulin locus SNPs in field populations of *T. circumcincta*. Red colour indicates F167Y (T<u>A</u>C), green indicates E198L (T<u>T</u>A) and blue indicates F200Y (T<u>A</u>C) and purple indicates susceptible alleles. Fig 4a: dots represent the resistance allele frequencies for each sampling time-point. The data shows the three farms with date of sample collection on the X-axis and the Y-axis representing the allele frequencies. Fig 4b: the pie charts show the overall prevalence of the isotype 1 β-tubulin locus SNPs in ewes and lambs on each farm. The prevalence of SNPs and combinations of SNPs at farm level are shown in the table below the figure.

In terms of overall prevalence, the F200Y (T<u>A</u>C) resistance-associated SNP was found on all three farms in each of the ewe and lamb samples collected at different time-points. The F167Y (T<u>A</u>C) resistance associated SNP was detected in 9/12 ewe and 7/13 lamb time-point samples on Farm 1; in 7/20 ewe and 7/9 lamb time-point samples on Farm 2; and in 2/11 ewe and 5/9 lamb time-point samples on Farm 3. The E198L (T<u>T</u>A) resistance associated SNP was detected in 12/12 ewe and 11/13 lamb time-point samples on Farm 1; 15/20 ewe and 9/9 lamb time-point samples on Farm 2; and 5/11 ewe and 8/9 lamb time-point samples on Farm 3.

## 4 Discussion

Genotypic markers and sensitive and accurate platforms with which to measure them are needed for accurate surveillance to assess and mitigate the impacts of climate, animal husbandry and pasture management on the emergence and spread of anthelmintic resistance mutations. Phenotypic methods of estimation of anthelmintic resistance are generally imprecise. *In vivo* bioassays such as the egg hatch (Le Jambre, 1976) and larval development (Coles et al., 1988) tests for benzimidazole resistance may be influenced by extrinsic or separate genetic factors governing traits such as fitness in the test environment, or by variation in the test conditions (Calvete et al., 2014). Furthermore, the mechanisms of drug resistance in eggs and developing larvae may differ from those in parasitic stages (Kotze et al., 2002). The *in vivo* faecal egg count reduction test is influenced by independent host factors, such as those affecting drug bioavailability (Ali and Hennessy, 1996), or by the accuracy or drug administration. Phenotypic assays are also influenced by the number of genes involved and by the dominance or recessiveness of the trait; hence are poor in the estimation of the frequency of resistance alleles, in particular when the frequency is low. Hence, they and may not accurately reflect the emergence and spread of anthelmintic resistance mutations in field populations of gastrointestinal nematodes.

Validated genotypic markers for anthelmintic resistance are only currently available for the benzimidazoles (Geary et al., 1992), albeit marker discovery is a research priority for the imidazothiazole (Sarai et al., 2013), macrocyclic lactone (Laing et al., 2013) and amino-acetonitrile derivative (Rufener et al., 2013) broad spectrum anthelmintic drug groups. In this study, we used the isotype 1 β-tubulin SNPs to develop a proof of concept method to examine the frequency of alleles in gastrointestinal nematode populations, with reference to better understanding of the impact of environmental factors on the emergence and spread of anthelmintic resistance. Deep amplicon sequencing by Illumina Mi-seq offers potential for greater practicality that is required for field investigations than pyrosequencing, because: fewer replicates are needed to overcome the chance of quality control failure; and separate pyrosequencing runs are required for analysis of each isotype 1 β-tubulin SNP, spanning codons 167, 198 and 200, whereas the Illumina Mi-seq platform allows for the determination of multi-allelic polymorphisms spanning an approximately 400 bp locus. Illumina Mi-seq involves two short PCRs and two rounds of product purification before sequencing, but in our hands provided informative read depths in at least 384 samples (four 96-well plates) in a single library; hence is most practical for use in high throughput scenarios.

We validated the Illumina Mi-seq method to examine benzimidazole resistance allele frequencies in mock population pools of *T. circumcincta* by: assessing sequence representation bias in the isotype 1 β-tubulin locus; comparing the results of Illumina Mi-seq and pyrosequencing; and applying the method to populations containing known proportions of resistant and susceptible L_3_. We identified no significant variation in the frequency of resistance alleles with the number of first round PCR cycles in any of the six reference *T. circumcincta* populations; hence showed no sequence representation bias (Avramenko et al., 2015) arising from the number of first round PCR cycles, potential variation in the efficiency of other library preparation steps, or acknowledged inaccuracy in the estimation due to random number of larvae (∼200 L_3_) making up each mock population pool. The Illumina Mi-seq method showed a higher F200Y (T<u>A</u>C) resistance allele frequency than pyrosequencing in each of the six reference *T. circumcincta* populations, but the differences were not shown to be statistically significant. This might imply that the Illumina Mi-seq method is more sensitive than pyrosequencing, but true determination of sensitivity would require the genotyping the individual L_3_ making up each of the mock population pools, which was beyond the limitations of this study. Frequencies of the F200Y (T<u>A</u>C) and F167Y (T<u>A</u>C) mutations of less than 5%, with low standard errors, were only detected by deep amplicon sequencing, implying that the method may overcome some of the weaknesses of pyrosequencing previously described in field studies of the emergence and spread of benzimidazole resistance mutations (Chaudhry et al., 2015). Five mock population pools containing different estimated proportions of homozygous F200Y (T<u>A</u>C) resistance mutations of were generated by mixing fixed amounts of pyrosequence genotyped individual L_3_ gDNA, derived from susceptible and drug selected resistant laboratory *T. circumcincta* populations. For the pools that were 100% resistant or 100% susceptible, the observed allele frequencies based on Illumina MiSeq were in perfect agreement with expected allele frequencies. For pools with 67%, 50%, and 33% resistance alleles, differences between the expected and observed frequencies of the F200Y (T<u>A</u>C) mutation were not statistically significant (p >0.9), with little variation between replicates, providing support for the accuracy of the Illumina Mi-seq method in determining the frequency of resistance alleles in a gastrointestinal nematode parasite population. The small amount of variation between observed and expected allele frequencies could have arisen in the creation of the mock pools using only 1μl of low concentration gDNA derived from individual L_3_. It is unlikely that this situation would be replicated in field samples comprised of large numbers of eggs or L_3_, hence having more representative mixing of alleles in gDNA.

Gastrointestinal nematodes impact heavily on animal welfare and production, hence there is a need to understand the population genetics of anthelmintic resistance as significant threat to global food security (Kaplan and Vidyashankar, 2012). Having validated the Illumina Mi-Seq platform using mock pools of laboratory *T. circumcincta* isolates, we applied the method to field samples collected from ewes and lambs on three farms, each highlighting different aspects of sheep management and approaches to parasite control. This was undertaken as proof of concept to explore possibilities for the application of a high throughput practical method to determine the proportions of resistance alleles in particular gastrointestinal nematode parasite species within mixed species field populations. The occurrence of multiple isotype 1 β-tubulin SNPs associated with the anthelmintic resistance phenotype in an individual gastrointestinal nematode is extremely rare (data on file), hence we examined the frequency of the total of the F200Y (T<u>A</u>C), E198L(<u>TT</u>C) and F167Y (T<u>A</u>C) mutations with reference to resistance allele frequencies in the field *T. circumcincta* populations. It has been suggested that the F167Y (T<u>A</u>C) and E198L (<u>TT</u>A) mutations cause resistance at lower doses of benzimidazole than the F200Y(T<u>A</u>C) mutation (Barrère et al., 2012), but this was discounted as a confounding factor in our study, because each mutation causes resistance at the recommended dose rates.

A different pattern of resistance allele frequency over time emerged on Farm 1 compared with Farm 2 and Farm 3. On farm 1, the overall frequencies of resistance alleles in ewes and lambs were about 94% and 90%, respectively, whereas the overall frequencies of resistance alleles in ewes and lambs on Farm 2/Farm 3 were 58%/70% and 77%/79%, respectively. There was less between sample variation on Farm 1 than on Farm 2 and Farm 3, some of which could have been stochastic, arising from low faecal egg counts and low proportions of *T. circumcincta* at the time of sampling. Together these results indicate high frequencies of benzimidazole resistance alleles on all three farms, having nearly reached genetic fixation on Farm 1. This scenario highlights an opportunity to use the results of high throughput anthelmintic resistance allele genotyping to investigate the genetic selection pressures and question management practices that might have impacted on them. For example, there had been little uptake of mitigation strategies on Farm 1 following the first diagnosis of anthelmintic resistance in 2000 (data on file); whereas Farm 2 had ceased anthelmintic treatments of periparturient ewes and implemented alternative grazing management with cattle following the diagnosis of anthelmintic resistance in 2001 (Sargison et al., 2001); and Farm 3 had cautiously reduced the anthelmintic treatment frequency of ewes and lambs and adopted refugia management strategies involving delayed treatments of lambs moved onto safe in-bye pastures, and avoidance of whole group treatments following the diagnosis of anthelmintic resistance in 2005 (Sargison et al., 2005). It is intriguing to consider the genetic fixation of benzimidazole resistance alleles on Farm 1 in the context of the sheep flock being open, having introduced replacement ewes annually that presumably harboured benzimidazole susceptible genotypes. The gene flow of alleles conferring susceptibility to benzimidazole drugs on Farm 1 may have been halted by strategic treatments of the introduced animals with amino-acetonitrile derivative, or anthelmintic drug combinations. While complicated by the need to mitigate the impact of resistance to each of the broad spectrum anthelmintic drug classes at once, this challenges the application of dogma surrounding quarantine treatments of introduced animals with anthelmintic drug combinations (Leathwick and Besier, 2014), without prior understanding of the population genetics of resistance alleles at an individual farm level. Once genetic fixation of resistance alleles has occurred, the probabilities of success of key mitigation strategies, such as the use of anthelmintic drug combinations (Leathwick et al., 2015), or refugia management involving targeted selective treatments (Greer et al., 2009), in achieving reversion to drug susceptibility are reduced.

On Farm 2 and Farm 3, where genetic fixation had not been reached, there was some between sample variations in the frequency of benzimidazole resistance alleles. Some of this would have been due to stochastic effects caused by different numbers of *T. circumcincta* L_3_ in the sample pools, but the observation nevertheless highlights opportunities to map sustained trends in the frequency of resistance alleles, indicating genetic drift or bottlenecks, to climatic variation and specific practices, such as anthelmintic drug treatments and grazing management. This was unrewarding in this case, where no benzimidazole treatments were administered in Farm 2 and Farm 3, and possibly because the frequency of resistance alleles was already too high to detect significant changes; showing a need for molecular surveillance starting before resistance reaches a level where it can be phenotypically identified. The higher frequency of resistance alleles in lambs compared to ewes in Flock 2 and Flock 3, raises questions about the relative selection pressures applied *T. circumcincta* in each group, implying that lamb treatments may have had the greatest impact.

In summary, there is a pressing need for high throughput methods to determine the frequency of anthelmintic resistance conferring mutations, or markers, in field populations of gastrointestinal nematodes. We have demonstrated the feasibility and practicality of deep amplicon sequencing by Illumina Mi-seq in this regard, using benzimidazole resistance in *T. circumcincta* as proof of concept, and have considered how knowledge of resistance allele frequencies in field populations might be used to build hypotheses on selection pressures and inform mitigation strategies. The method could be applied to other gastrointestinal nematode species, and multiplexed to study benzimidazole resistance co-infections (Avramenko et al., 2019). Once markers become available, the method might also be used to study single or multiple mutations conferring resistance to other anthelmintic drug groups, and combined with phylogenetic tools to show their emergence and spread.

