## Supplementary Table S1 for "Development of high throughput method for the analysis of anthelmintic resistance allele frequencies in field populations of gastrointestinal nematodes"

**Supplementary Table S1**: Primer sequences used for this study. Presented results of pyrosequencing used the P167 TC, P198 TC, and P200 TC primers. Illumina adapters are the Nextera transposase adapters, and for each direction, there are four versions with a different number of random nucleotides (N).

| Primer | Purpose | Sequence, 5’-3’ | Reference |
| --- | --- | --- | --- |
| BTUB_FOR | Amplifying β-tubulin for pyrosequencing | CCAAAATTCGCGAGGAGTA | Skuce *et al*. 2010 |
| BTUB_REV (biotin) | Amplifying β-tubulin for pyrosequencing | 5Bioag/TTTCAAGGTGCGGAAGCAGA |  |
| P167 TC | Pyrosequencing | ATAGAATCATGGCTTCAT | Unpublished |
| P198 TC | Pyrosequencing | GGTWGAAAAYACCGAYK |  |
| P200 TC | Pyrosequencing | GAAAAYACCGATGAAACRT |  |
| BTUB_FOR with Illumina adapter | Amplifying β-tubulin for Illumina Mi-seq | TCGTCGGCAGCGTCAGATGTGTATAAGAGACAGCCAAAATTCGCGAGGAG*T*A | Oligonucleotide sequences © 2018 Illumina, Inc. All rights reserved |
| BTUB_FOR with Illumina adapter (1N) | Amplifying β-tubulin for Illumina Mi-seq | TCGTCGGCAGCGTCAGATGTGTATAAGAGACAGNCCAAAATTCGCGAGGAG*T*A |  |
| BTUB_FOR with Illumina adapter (2N) | Amplifying β-tubulin for Illumina Mi-seq | TCGTCGGCAGCGTCAGATGTGTATAAGAGACAGNNCCAAAATTCGCGAGGAG*T*A |  |
| BTUB_FOR with Illumina adapter (3N) | Amplifying β-tubulin for Illumina Mi-seq | TCGTCGGCAGCGTCAGATGTGTATAAGAGACAGNNNCCAAAATTCGCGAGGAG*T*A |  |
| BTUB_REV with Illumina adapter | Amplifying β-tubulin for Illumina Mi-seq | GTCTCGTGGGCTCGGAGATGTGTATAAGAGACAGTTTCAAGGTGCGGAAGCA*G*A |  |
| BTUB_REV with Illumina adapter (1N) | Amplifying β-tubulin for Illumina Mi-seq | GTCTCGTGGGCTCGGAGATGTGTATAAGAGACAGNTTTCAAGGTGCGGAAGCA*G*A |  |
| BTUB_REV with Illumina adapter (2N) | Amplifying β-tubulin for Illumina Mi-seq | GTCTCGTGGGCTCGGAGATGTGTATAAGAGACAGNNTTTCAAGGTGCGGAAGCA*G*A |  |
| BTUB_REV with Illumina adapter (3N) | Amplifying β-tubulin for Illumina Mi-seq | GTCTCGTGGGCTCGGAGATGTGTATAAGAGACAGNNNTTTCAAGGTGCGGAAGCA*G*A |  |
