## Supplementary Table S3 for "Development of high throughput method for the analysis of anthelmintic resistance allele frequencies in field populations of gastrointestinal nematodes"

**Supplementary Table S3:** Mean benzimidazole resistance allele frequencies in pools from six *T. circumcincta* laboratory isolates amplified at different numbers of PCR cycles (25X, 30X, 35X, 40X). The F200Y (TAC) mutations were identified in all six populations and F167Y (TAC) mutation detected in MTci13.

| **Sample** | **No. of PCR cycles** | **Mean no. of Illumina MiSeq reads** | **Mean no. of susceptible reads (Illumina MiSeq)** | **Mean no. of resistant reads (Illumina MiSeq)** | **F167Y (%)** | | **F200Y (%)** | |
| --- | --- | --- | --- | --- | --- | --- | --- | --- |
|  |  |  |  |  | TTC | TAC | TTC | TAC |
| MTci2 | 25X | 9458 | 8548 | 910 | 100.00 | 0.00 | 90.45 | 9.55 |
|  | 30X | 8754 | 7790 | 964 | 100.00 | 0.00 | 88.42 | 11.58 |
|  | 35X | 7842 | 6902 | 940 | 100.00 | 0.00 | 88.69 | 11.31 |
|  | 40X | 7990 | 7190 | 800 | 100.00 | 0.00 | 89.61 | 10.39 |
| MTci5 | 25X | 13917 | 4539 | 9378 | 100.00 | 0.00 | 32.64 | 67.36 |
|  | 30X | 11952 | 3765 | 8187 | 100.00 | 0.00 | 31.49 | 68.51 |
|  | 35X | 10397 | 3270 | 7127 | 100.00 | 0.00 | 31.82 | 68.18 |
|  | 40X | 12987 | 4073 | 8914 | 100.00 | 0.00 | 31.90 | 68.10 |
| MTci7 | 25X | 13099 | 3227 | 9872 | 100.00 | 0.00 | 24.30 | 75.70 |
|  | 30X | 9852 | 2495 | 7357 | 100.00 | 0.00 | 25.22 | 74.78 |
|  | 35X | 12556 | 2938 | 9618 | 100.00 | 0.00 | 23.46 | 76.54 |
|  | 40X | 12552 | 2815 | 9737 | 100.00 | 0.00 | 22.72 | 77.28 |
| MTci11 | 25X | 9601 | 9304 | 297 | 100.00 | 0.00 | 96.91 | 3.39 |
|  | 30X | 4186 | 4015 | 171 | 100.00 | 0.00 | 95.91 | 4.16 |
|  | 35X | 4124 | 3980 | 144 | 100.00 | 0.00 | 96.51 | 3.70 |
|  | 40X | 5419 | 5210 | 209 | 100.00 | 0.00 | 96.14 | 3.01 |
| MTci12 | 25X | 4021 | 309 | 3712 | 100.00 | 0.00 | 7.83 | 92.17 |
|  | 30X | 8850 | 653 | 8197 | 100.00 | 0.00 | 7.52 | 92.48 |
|  | 35X | 5602 | 393 | 5209 | 100.00 | 0.00 | 7.23 | 92.77 |
|  | 40X | 5681 | 306 | 5375 | 100.00 | 0.00 | 5.50 | 94.50 |
| MTci13 | 25X | 14757 | 4900 | 9857 | 97.38 | 2.62 | 33.57 | 66.43 |
|  | 30X | 11120 | 3422 | 7698 | 97.20 | 2.80 | 30.29 | 69.71 |
|  | 35X | 11113 | 3577 | 7536 | 95.72 | 4.28 | 32.85 | 67.15 |
|  | 40X | 9640 | 2947 | 6693 | 95.88 | 4.12 | 30.57 | 69.43 |
