## Supplementary Table S4 for "Development of high throughput method for the analysis of anthelmintic resistance allele frequencies in field populations of gastrointestinal nematodes"

**Supplementary Table S4:** Benzimidazole resistance allele frequencies in pools of *T. circumcincta* laboratory isolates with about 200 L_3_, as determined by two methods: Illumina Mi-seq and pyrosequencing genotyping. Both F167Y (TAC) and F200Y (TAC) mutations were identified by Illumina Mi-seq and only F200Y (TAC) mutation was found by pyrosequencing.

| **Method** | **Sample** | **Mean no. of Illumina MiSeq reads** | **Mean no. of susceptible reads (Illumina MiSeq)** | **Mean no. of resistant reads (Illumina MiSeq** | **F167Y (%)** | | **F200Y (%)** | |
| --- | --- | --- | --- | --- | --- | --- | --- | --- |
|  |  |  |  |  | TTC | TAC | TTC | TAC |
| **Illumina MiSeq** | MTci2 | 7842 | 6902 | 940 | 100.00 | 0.00 | 88.69 | 11.31 |
|  | MTci5 | 10397 | 3270 | 7127 | 100.00 | 0.00 | 31.82 | 68.18 |
|  | MTci7 | 12556 | 2938 | 9618 | 100.00 | 0.00 | 23.46 | 76.54 |
|  | MTci11 | 4124 | 3980 | 144 | 100.00 | 0.00 | 96.51 | 3.70 |
|  | MTci12 | 5602 | 393 | 5209 | 100.00 | 0.00 | 7.23 | 92.77 |
|  | MTci13 | 11113 | 3577 | 7536 | 95.72 | 4.28 | 32.85 | 67.15 |
| **Pyrosequencing** | MTci2 | Not applicable | | | 0.0 | 100.0 | 91.8 | 8.2 |
|  | MTci5 |  |  |  | 0.0 | 100.0 | 41.5 | 58.5 |
|  | MTci7 |  |  |  | 0.0 | 100.0 | 27.2 | 72.8 |
|  | MTci11 |  |  |  | 0.0 | 100.0 | 100.0 | 0.0 |
|  | MTci12 |  |  |  | 0.0 | 100.0 | 14.7 | 85.3 |
|  | MTci13 |  |  |  | 0.0 | 100.0 | 34.3 | 65.7 |
