## Supplementary Table S5 for "Development of high throughput method for the analysis of anthelmintic resistance allele frequencies in field populations of gastrointestinal nematodes"

**Supplementary Table S5:** Mean frequency of the F200Y (TAC) mutation for benzimidazole resistance, made using pyrosequence genotyped individual larvae from phenotypically benzimidazole susceptible (MTci2) and resistant (MTci12) *T. circumcincta* laboratory populations*.* The expected frequency is calculated based on how the pools were made.

| **Pooled samples** | **Mean no. of Illumina MiSeq reads** | **Mean no. of susceptible reads (Illumina MiSeq)** | **Mean no. of resistant reads (Illumina MiSeq)** | **Observed**  **frequency**  **for F200Y (%)** | | | | **Expected**  **frequency**  **for F200Y (%)** | |
| --- | --- | --- | --- | --- | --- | --- | --- | --- | --- |
|  |  |  |  | | TTC | TAC | TTC | | TAC |
| MixS | 7195 | 7195 | 0 | | 100.0 | 0.0 | 100 | | 0 |
| MixR | 8147 | 0 | 8147 | | 0.0 | 100.0 | 0 | | 100 |
| MixSR | 8452 | 5731 | 2721 | | 67.8 | 32.2 | 50 | | 50 |
| MixSRR | 9302 | 2006 | 7296 | | 21.6 | 78.4 | 33 | | 66 |
| MixSSR | 7943 | 6103 | 1840 | | 76.8 | 23.2 | 66 | | 33 |
