## Supplementary Table S6 for "Development of high throughput method for the analysis of anthelmintic resistance allele frequencies in field populations of gastrointestinal nematodes"

**Supplementary Table S6:** Benzimidazole resistance allele frequency of field samples taken from three farms in south-east Scotland from ewes and lambs, monthly over the period of 2016-2017. The F167Y (TAC), E198L (TTA), and F200Y (TAC) mutations are shown.

| **Farm** | **Sheep** | **Date** | **Total no. of Illumina MiSeq reads** | **Total no. of susceptible reads (Illumina MiSeq)** | **Total no. of resistant reads (Illumina MiSeq)** | **F167Y (%)** | | **E198L (%)** | | **F200Y (%)** | |
| --- | --- | --- | --- | --- | --- | --- | --- | --- | --- | --- | --- |
|  |  |  |  |  |  | TTC | TAC | GAA | TTA | TTC | TAC |
| **Farm 1** | ewe | 20/3/16 | 74093 | 3504 | 70589 | 99.1 | 0.9 | 96.8 | 3.2 | 8.8 | 91.2 |
|  |  | 7/4/16 | 155344 | 10619 | 144725 | 99.2 | 0.8 | 97.2 | 2.8 | 10.5 | 89.5 |
|  |  | 30/5/16 | 12562 | 324 | 12238 | 93.4 | 6.6 | 96.6 | 3.4 | 12.6 | 87.4 |
|  |  | 7/6/16 | 1271 | 25 | 1246 | 100 | 0 | 95.9 | 4.1 | 6.0 | 94.0 |
|  |  | 10/8/16 | 8884 | 1364 | 7520 | 100 | 0 | 97.0 | 3.0 | 18.3 | 81.7 |
|  |  | 13/10/16 | 23804 | 196 | 23608 | 99.3 | 0.7 | 95.7 | 4.3 | 5.8 | 94.2 |
|  |  | 23/11/16 | 121067 | 9123 | 111944 | 98.8 | 1.2 | 95.9 | 4.1 | 12.9 | 87.1 |
|  |  | 20/2/17 | 9078 | 303 | 8775 | 99.0 | 1.0 | 97.3 | 2.7 | 7.0 | 93.0 |
|  |  | 16/3/17 | 44947 | 4744 | 40203 | 98.4 | 1.6 | 99.4 | 0.6 | 12.7 | 87.3 |
|  |  | 17/4/17 | 44393 | 1607 | 42786 | 98.9 | 1.1 | 93.8 | 6.2 | 10.9 | 89.1 |
|  |  | 18/4/17 | 80316 | 2983 | 77333 | 99.5 | 0.5 | 96.5 | 3.5 | 7.7 | 92.3 |
|  |  | 21/5/17 | 41100 | 6542 | 34558 | 100 | 0 | 94.2 | 5.8 | 21.7 | 78.3 |
|  | lamb | 7/6/16b | 129105 | 1165 | 127940 | 98.2 | 1.8 | 100 | 0 | 2.7 | 97.3 |
|  |  | 18/7/16 | 13202 | 0 | 13202 | 100 | 0 | 90.3 | 9.7 | 9.7 | 90.3 |
|  |  | 8/8/16 | 74071 | 13361 | 60710 | 100 | 0 | 97.4 | 2.6 | 20.6 | 79.4 |
|  |  | 1/9/16 | 673 | 81 | 592 | 99.0 | 1.0 | 99.3 | 0.7 | 13.8 | 86.2 |
|  |  | 20/9/16 | 13075 | 455 | 12620 | 98.4 | 1.6 | 95.6 | 4.4 | 9.5 | 90.5 |
|  |  | 20/9/16b | 1101 | 53 | 1048 | 99.0 | 1.0 | 97.2 | 2.8 | 8.6 | 91.4 |
|  |  | 27/9/16 | 138303 | 5505 | 132798 | 99.4 | 0.6 | 96.7 | 3.3 | 7.9 | 92.1 |
|  |  | 1/11/16 | 167890 | 29691 | 138199 | 100 | 0 | 96.4 | 3.6 | 21.3 | 78.7 |
|  |  | 1/11/16b | 8902 | 1 | 8901 | 100 | 0 | 100 | 0 | 0.0 | 100.0 |
|  |  | 23/11/16b | 228183 | 12483 | 215700 | 97.5 | 2.5 | 98.1 | 1.9 | 9.9 | 90.1 |
|  |  | 14/6/17 | 32402 | 11685 | 20717 | 99.4 | 0.6 | 84.1 | 15.9 | 52.5 | 47.5 |
|  |  | 10/8/17 | 2548 | 345 | 2203 | 100 | 0 | 98.2 | 1.8 | 15.3 | 84.7 |
|  |  | 30/8/17 | 29910 | 4268 | 25642 | 100 | 0 | 95.8 | 4.3 | 18.5 | 81.5 |
| **Farm 2** | ewe | 29/3/16 | 2133 | 802 | 1331 | 100 | 0 | 98.1 | 1.9 | 39.5 | 60.5 |
|  |  | 18/4/16 | 11905 | 3734 | 8171 | 98.8 | 1.2 | 99.1 | 0.9 | 33.4 | 66.6 |
|  |  | 27/5/16 | 67572 | 29276 | 38296 | 96.2 | 3.8 | 100 | 0 | 47.2 | 52.8 |
|  |  | 14/7/16d | 34037 | 8373 | 25664 | 100 | 0 | 98.8 | 1.2 | 25.8 | 74.2 |
|  |  | 26/7/16 | 1608 | 354 | 1254 | 100 | 0 | 96.2 | 3.8 | 25.7 | 74.3 |
|  |  | 26/8/16 | 13921 | 4375 | 9546 | 96.6 | 3.4 | 97.8 | 2.2 | 37.1 | 62.9 |
|  |  | 7/9/16 | 32373 | 9316 | 23057 | 100 | 0 | 98.8 | 1.2 | 30 | 70 |
|  |  | 20/9/16d | 12789 | 8602 | 4187 | 100 | 0 | 86.1 | 13.9 | 81.2 | 18.8 |
|  |  | 17/10/16 | 43935 | 39501 | 4434 | 100 | 0 | 100 | 0 | 89.9 | 10.1 |
|  |  | 26/2/17b | 29809 | 8932 | 20877 | 100 | 0 | 99.2 | 0.8 | 30.8 | 69.2 |
|  |  | 22/3/17 | 25695 | 7201 | 18494 | 98.8 | 1.2 | 99.3 | 0.7 | 30.0 | 70.0 |
|  |  | 5/4/17 | 24862 | 8409 | 16453 | 98.9 | 1.1 | 99.9 | 0.1 | 35.0 | 65.0 |
|  |  | 17/4/17b | 64038 | 28315 | 35723 | 100 | 0 | 100 | 0 | 44.2 | 55.8 |
|  |  | 17/4/17c | 18779 | 8142 | 10637 | 100 | 0 | 99.99 | 0.01 | 43.4 | 56.6 |
|  |  | 3/5/17 | 23050 | 23048 | 2 | 100 | 0 | 100 | 0 | 99.99 | 0.01 |
|  |  | 30/5/17 | 1378 | 404 | 974 | 99.6 | 0.4 | 91.1 | 8.9 | 38.7 | 61.3 |
|  |  | 30/5/17b | 35176 | 5694 | 29482 | 99.4 | 0.6 | 99.6 | 0.4 | 17.2 | 82.8 |
|  |  | 14/6/17b | 19288 | 741 | 18547 | 100 | 0 | 96.4 | 3.6 | 7.4 | 92.6 |
|  |  | 14/6/17c | 1091 | 832 | 259 | 100 | 0 | 99.99 | 0.01 | 76.2 | 23.8 |
|  |  | 4/7/17 | 46160 | 22970 | 23190 | 100 | 0 | 100 | 0 | 49.8 | 50.2 |
|  | lamb | 26/7/16b | 108013 | 18957 | 89056 | 98.1 | 1.9 | 97.2 | 2.8 | 22.3 | 77.7 |
|  |  | 5/8/16 | 59282 | 10578 | 48704 | 98.7 | 1.3 | 99.3 | 0.7 | 19.8 | 80.2 |
|  |  | 6/8/16 | 110047 | 14776 | 95271 | 96.6 | 3.4 | 92.8 | 7.2 | 24.0 | 76.0 |
|  |  | 6/8/16b | 55742 | 9183 | 46559 | 100 | 0 | 93.8 | 6.2 | 22.7 | 77.3 |
|  |  | 19/8/16 | 17509 | 3101 | 14408 | 98.9 | 1.1 | 99.0 | 1.0 | 19.8 | 80.2 |
|  |  | 7/9/16b | 6092 | 1545 | 4547 | 99.0 | 1.0 | 99.2 | 0.8 | 27.1 | 72.9 |
|  |  | 20/9/16e | 15558 | 4298 | 11260 | 100 | 0 | 96.8 | 3.2 | 30.9 | 69.1 |
|  |  | 17/10/16b | 20147 | 7062 | 13085 | 99.1 | 0.9 | 97.7 | 2.3 | 38.3 | 61.7 |
|  |  | 24/11/16 | 78589 | 26699 | 51890 | 98.2 | 1.8 | 99.9 | 0.1 | 35.9 | 64.1 |
| **Farm 3** | ewe | 18/2/16 | 18607 | 6456 | 12151 | 100 | 0 | 100 | 0 | 34.7 | 65.3 |
|  |  | 18/3/16 | 41362 | 5640 | 41361 | 100 | 0 | 0 | 0 | 14.5 | 85.5 |
|  |  | 24/5/16 | 18833 | 2177 | 16656 | 100 | 0 | 99.9 | 0.1 | 11.6 | 88.4 |
|  |  | 3/6/16 | 36011 | 10887 | 25124 | 100 | 0 | 100 | 0 | 30.2 | 69.8 |
|  |  | 23/6/16 | 3052 | 1032 | 2020 | 100 | 0 | 100 | 0 | 33.8 | 66.2 |
|  |  | 14/7/16 | 7274 | 1384 | 5890 | 100 | 0 | 99.99 | 0.01 | 19.0 | 81.0 |
|  |  | 17/8/16 | 12664 | 9458 | 3206 | 100 | 0 | 100 | 0 | 74.7 | 25.3 |
|  |  | 12/9/16 | 34121 | 8613 | 25508 | 97.8 | 2.2 | 99.9 | 0.1 | 27.6 | 72.4 |
|  |  | 26/10/16 | 16020 | 2572 | 13448 | 97.7 | 2.3 | 94.2 | 5.8 | 24.2 | 75.8 |
|  |  | 26/2/17 | 15026 | 5061 | 9965 | 100 | 0 | 100 | 0 | 33.7 | 66.3 |
|  |  | 20/3/17 | 47673 | 26491 | 21182 | 100 | 0 | 86.3 | 13.7 | 69.3 | 30.7 |
|  | lamb | 30/6/16 | 70399 | 20735 | 49664 | 100 | 0 | 94.0 | 6.0 | 35.5 | 64.6 |
|  |  | 14/7/16b | 38869 | 5384 | 33485 | 97.4 | 2.6 | 90.8 | 9.2 | 25.7 | 74.3 |
|  |  | 2/8/16 | 71590 | 8492 | 63098 | 98.4 | 1.6 | 96.2 | 3.8 | 17.3 | 82.7 |
|  |  | 17/8/16b | 58397 | 17401 | 40996 | 100 | 0 | 96.1 | 3.9 | 33.7 | 66.3 |
|  |  | 12/9/16b | 61649 | 7437 | 54212 | 97.0 | 3.0 | 91.5 | 8.5 | 23.6 | 76.4 |
|  |  | 6/10/16 | 43384 | 12411 | 30973 | 97.6 | 2.4 | 99.7 | 0.3 | 31.4 | 68.6 |
|  |  | 20/10/16 | 56366 | 17395 | 38971 | 100 | 0 | 100 | 0 | 30.9 | 69.1 |
|  |  | 14/11/16 | 77540 | 23251 | 54289 | 96.4 | 3.6 | 98.9 | 1.1 | 34.7 | 65.3 |
|  |  | 8/12/16 | 48489 | 3233 | 45256 | 100 | 0 | 96.9 | 3.1 | 9.8 | 90.2 |
